# Disrupting actin filaments enhances glucose-stimulated insulin secretion independent of the cortical actin cytoskeleton

**DOI:** 10.1101/2023.07.15.549141

**Authors:** Alexander J. Polino, Xue Wen Ng, Rebecca Rooks, David W. Piston

## Abstract

Just under the plasma membrane of most animal cells lies a dense meshwork of actin filaments called the cortical cytoskeleton. In insulin-secreting pancreatic β cells, a longstanding model posits that the cortical actin layer primarily acts to restrict access of insulin granules to the plasma membrane. Here we test this model and find that stimulating β cells with pro-secretory stimuli (glucose and/or KCl) has little impact on the cortical actin layer. Chemical perturbations of actin polymerization, by either disrupting or enhancing filamentation, dramatically enhances glucose-stimulated insulin secretion. We find that this enhancement does not correlate with the state of the cortical actin layer, suggesting filament disruptors act on insulin secretion independently of the cortical cytoskeleton.

## Introduction

Most animal cells are enveloped in a dense layer of actin filaments, called the cortical cytoskeleton. In coordination with myosins and other actin-binding proteins, this layer is dynamically remodeled, driving cell shape changes required for motility, division, and contractile tissue motion [1]. In models of regulated secretion, the cortex has been implicated in two opposing functions. First, the dense cortical network restricts access of secretory granules to the plasma membrane. Second, cortex remodeling enables secretion by providing a path to secretory sites and generating force to complete granule fusion with the plasma membrane [2,3].

In pancreatic β cells, cortical actin has been hypothesized to primarily restrict the access of insulin granules to the plasma membrane, limiting insulin secretion. This model rests on the observations that stimulating β cells with pro-secretory levels of glucose dramatically reduces levels of total cellular actin [4–8], and that forcing the depolymerization of the actin cortex with chemical disruptors enhances glucose-stimulated insulin secretion (GSIS) [9–11]. The first finding has been shown in immortalized β cell models INS-1 [4] and MIN6 [5], as well as in *ex vivo* murine islets [8]. However, various reports have found inconsistent results, ranging from no change in actin staining to over 50 percent loss of actin staining. The second finding was originally shown in Lelio Orci’s 1972 electron microscopy study describing cortical microfilaments in the β cells of rat islets; a 90-minute treatment with the actin depolymerizer cytochalasin B enhanced GSIS from rat islets by around 50% [10]. This finding has since been recapitulated in dispersed and intact mouse islets with a variety of chemicals and concentrations [8,9,11]. Curiously, Mourad, et al. (2013) found that pretreatment with the filament polymerizing drug jasplakinolide also enhances GSIS, phenocopying filament disruptors. This finding seems at odds with the model of cortical actin restricting granule secretion.

Here, we sought to test the model of cortical actin restricting insulin secretion by investigating each of its supporting observations with fluorescence and scanning electron microscopy, correlated with secretion assays. We find that glucose or KCl stimulation has only minimal impact on cortical actin filaments, both in MIN-6 cells and mouse pancreatic islets. Further, we find that modulating the actin filament structure by chemical inhibition of filament formation or stabilization of actin polymerization both dramatically enhance glucose-stimulated insulin secretion. Thus, the state of the cortical cytoskeleton does not explain the shared phenotype of actin disruption enhancing insulin secretion.

Instead, our data support a model where the effect of actin filament disruption on insulin secretion is independent of the cortical cytoskeleton.

## Results

### Stimulating β cells does not substantially change actin levels

We sought to expand upon prior observations by investigating actin levels at various times after stimulating β cells with glucose or KCl. Throughout this work, 11 mM glucose is used to stimulate *ex vivo* mouse islets; 20 mM glucose to stimulate the immortalized murine β cell model MIN6. 30 mM KCl is used as an alternative secretagogue, as it directly depolarizes the plasma membrane, resulting in secretion independent of the canonical glucose sensing pathway. To quantify F-actin levels, we stimulated MIN6 cells or murine islets with high glucose or high glucose plus KCl. We then fixed the samples at 0, 2, 5, 10, or 15 minutes after stimulation, stained the F-actin with phalloidin-AlexaFluor488, and images at the cell-coverslip boundary, where actin networks are most prominent. F-actin levels were quantified per cell area (Fig. 1A, B) or per islet area (Fig. 1C, D). We find only slight changes in global actin levels in stimulated MIN6 cells (one-way ANOVA, F(8, 149) = 2.138, p = 0.0356). Most group means vary by just a few percent from the unstimulated control. Dunnett’s multiple comparisons test shows that the only statistically significant difference is between control and 10-minute glucose + KCl treatment (adjusted p-value of 0.0109, and a 95% confidence interval of the magnitude of difference ranging from 0.02150 to 0.2384). Murine islets have more natural variation in size and actin staining than MIN6 cells, but still fail to show a consistent effect of pro-secretory stimuli on actin levels (one-way ANOVA, F(8, 130) = 3.021, p = 0.0038). Again, Dunnett’s multiple comparisons indicates that the only statistically significant difference is between control and 10-minute glucose treatment (adjusted p-value of 0.0067, and a 95% confidence interval of the difference ranging from 0.067 to 0.602). Given that the direction of change is opposite that seen in the MIN6 cells, not seen at other time points or in KCl-treated islets, and opposite of that described in prior literature, we suspect it is just an outlier as a result of biological variation of *ex vivo* islets, and not physiologically relevant.

**Figure 1.**
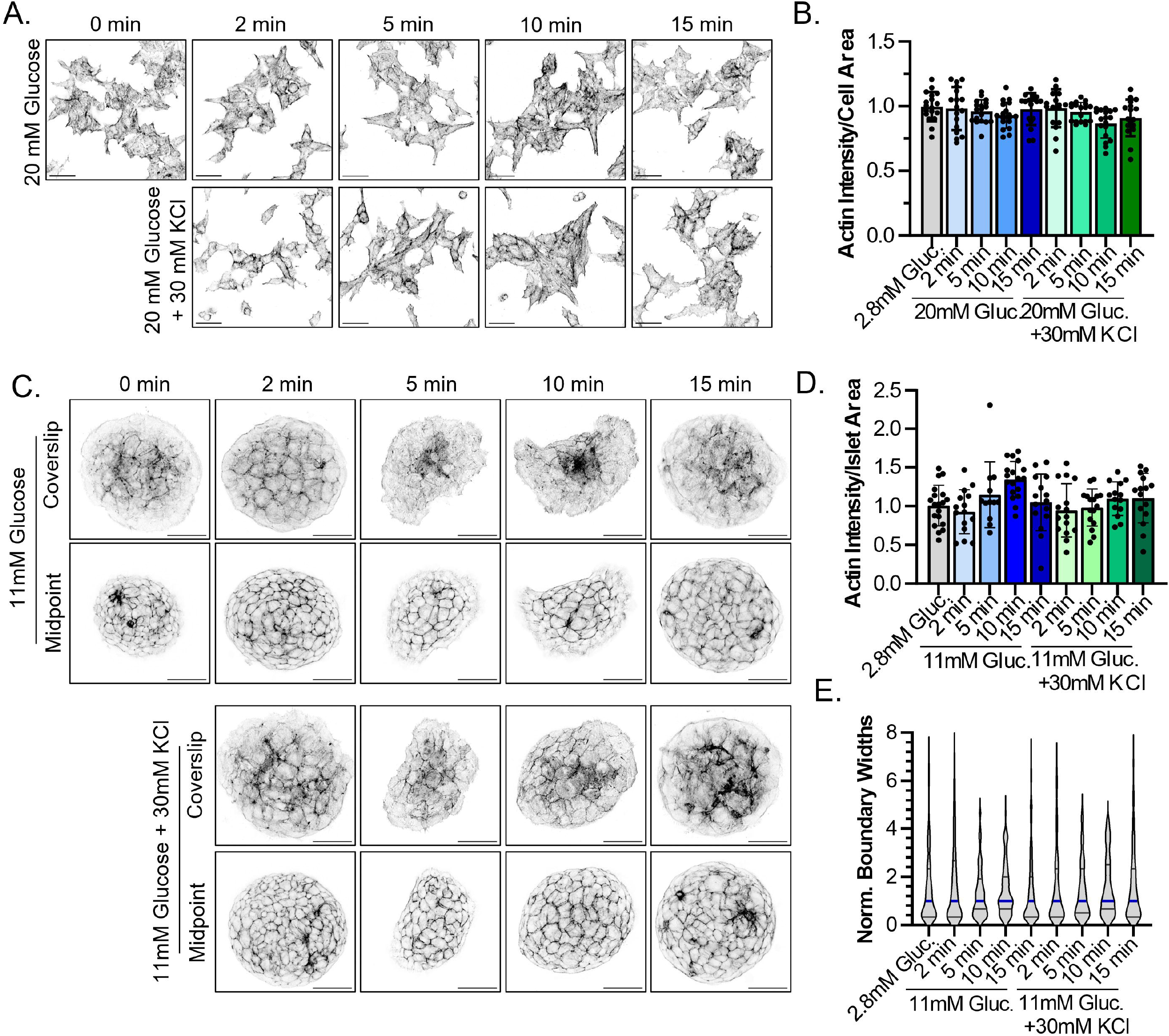
Stimulating β cells does not substantially change actin levels. (A) MIN6 cells were stimulated with high glucose (20 mM) +/-30mM KCl and fixed at 0, 2, 5, 10, or 15 min. Actin in fixed cells was stained with phalloidin-AlexaFluor488, and imaged. Representative images are shown. Scale bars represent 20 μm. (B) Graph comparing the actin intensity per cell area for each image. The experiment was performed three times, with six images quantified per condition in each experiment. All points are shown, each representing the actin intensity for a given image; error bars represent standard deviation. (C) Islets stimulated as in (A) with high glucose (11 mM) +/-30 mM KCl and fixed at times indicated. Each islet was imaged at the coverslip (upper row) to highlight actin networks at the coverslip interface, and at its midpoint (lower row) to highlight actin at cell-cell boundaries. Scale bars represent 40 μm. (D) Actin intensity was quantified per islet area. Experiment was performed three times with up to six islets imaged each time. All points are shown, each representing the actin intensity divided by a given islet’s area; error bars represent standard deviation. (E) The breadth of actin staining at cell-cell boundaries was measured from two perpendicular lines through five islets per condition. The thick blue line represents the median of the measured boundaries, the thin black lines the first and third quartiles.

We also quantified the F-actin signal by acquiring confocal images through the middle of each islet (Fig. 1C, “Midpoint”) where phalloidin signal corresponds primarily to the cortical actin ring around each cell. We measured the widths of these cell-cell boundaries, which again exhibit no substantial differences between treatment groups (Fig. 1E, Kruskal-Wallis test, p = 0.6682). As a verification of the fluorescence measurements of F-actin, we also quantified actin filamentation biochemically in MIN6 following stimulation for five minutes, as shown to stimulate actin rearrangements in prior reports [5,8]. We lysed the cells and separated intact actin filaments by ultracentrifugation. We detect no effect of glucose +/-KCl on global actin filament state (Fig. S1).

Cortical actin is a meshwork of filaments, with filaments too densely packed to be resolved by light microscopy. To image directly the effects of glucose/KCl stimulation on finer cortex architecture, we fixed MIN6 cells or mouse islets, detergent-extracted the plasma membranes, and used scanning electron microscopy (SEM) to visualize the cytoskeleton (Fig. 2). Consistent with the fluorescence and biochemical assays, we see no global change in cortical cytoskeleton arrangement after pro-secretory stimulation either in MIN6 cells (Fig. 2A, B) or islets (Fig. 2C, D). Quantifying the percent of the cell surface covered by filaments reveals no change with stimulation in MIN6 cells (one-way ANOVA, F(2, 33) = 1.215, p = 0.3097), and a slight increase in coverage with islet stimulation (one-way ANOVA, F(2, 51) = 22.64, p < 0.0001) of a magnitude less than 10% (Fig. 2B, D).

**Figure 2.**
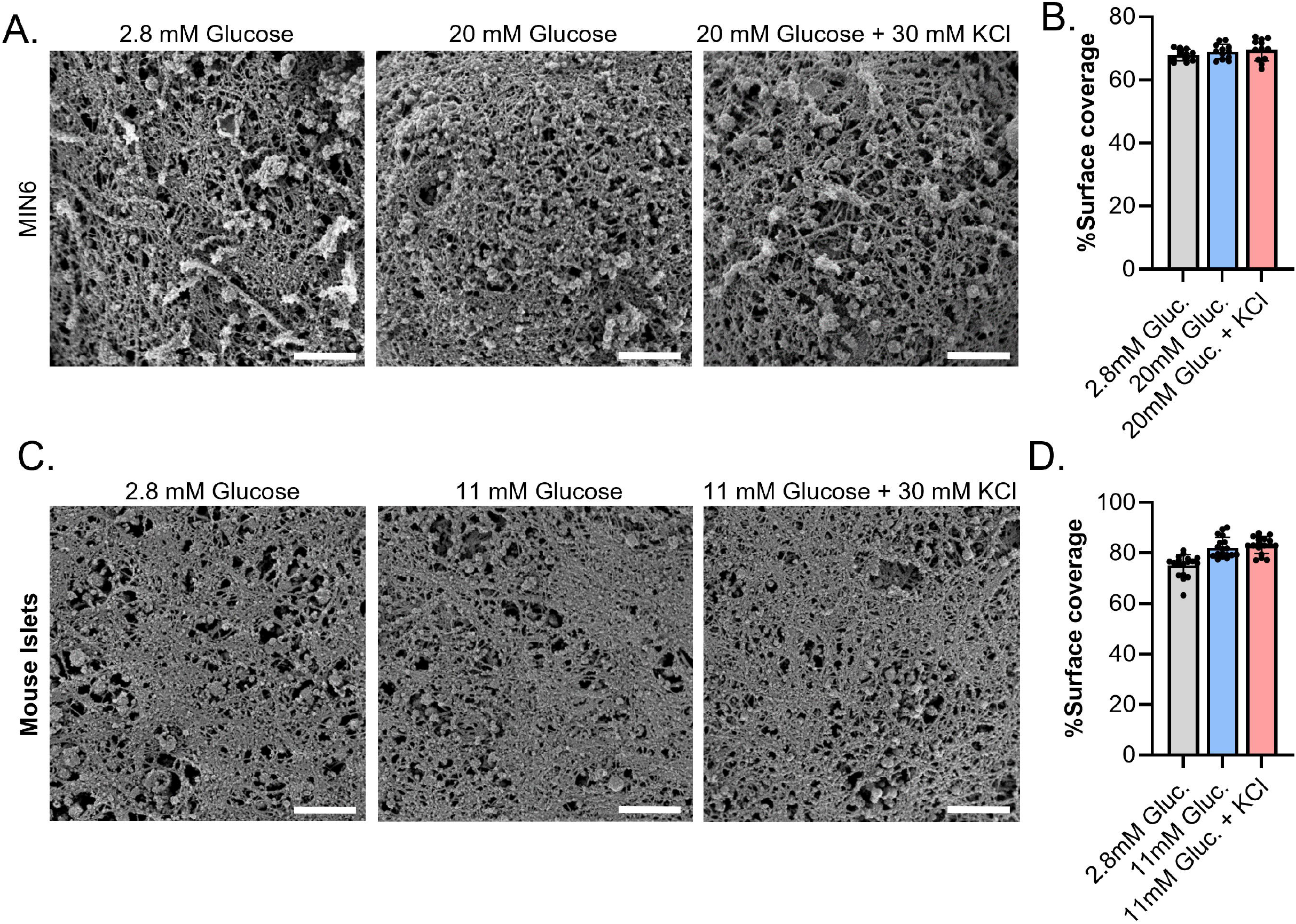
Stimulating β cells does not dramatically change the actin cortex. (A) SEM images of membrane-stripped MIN6 cells revealing the cortical cytoskeleton. Cells were treated with 2.8 mM glucose, 20 mM glucose, or 20 mM glucose + 30 mM KCl for 5 minutes prior to fixation and membrane stripping. Scale bars represent 1 μm. (B) The percent of the surface covered by cytoskeleton was quantified in 10-12 images per condition. Error bars represent standard deviations. (C) Mouse islets treated as above, but with 2.8 mM glucose, 11 mM glucose, or 11 mM glucose + 30 mM KCl. Scale bars represent 1 μm. (D) Six images from each of three islets were quantified per condition, as in (B). Error bars represent standard deviation. This experiment was performed twice with similar results. The results from one experiment are shown above. Zoomed out images are provided in Figure S2.

### Actin filament-disrupting drugs enhance GSIS

We next sought to expand upon the previous observation that actin depolymerizing agents enhance GSIS. We recapitulate this observation, finding that a 30-minute pre-treatment with actin depolymerizers enhances GSIS in a concentration-dependent manner. This is true for filament barbed-end binders cytochalasin B and D (Fig. 3A, B), filament pointed-end binders latrunculins A and B (Fig. 3C, D), and the filament-severing drug swinholide A (Fig. 3E). This is also consistent across both MIN6 cells (Fig. 3A-E), and mouse islets (Fig. 3F). Surprisingly, we see a similar dose-dependent enhancement of GSIS in cells or islets pre-treated with the filament enhancing drug jasplakinolide (Fig. 4). This raises the obvious question: why do compounds with seemingly opposite effects on actin have the same effect on insulin secretion?

**Figure 3.**
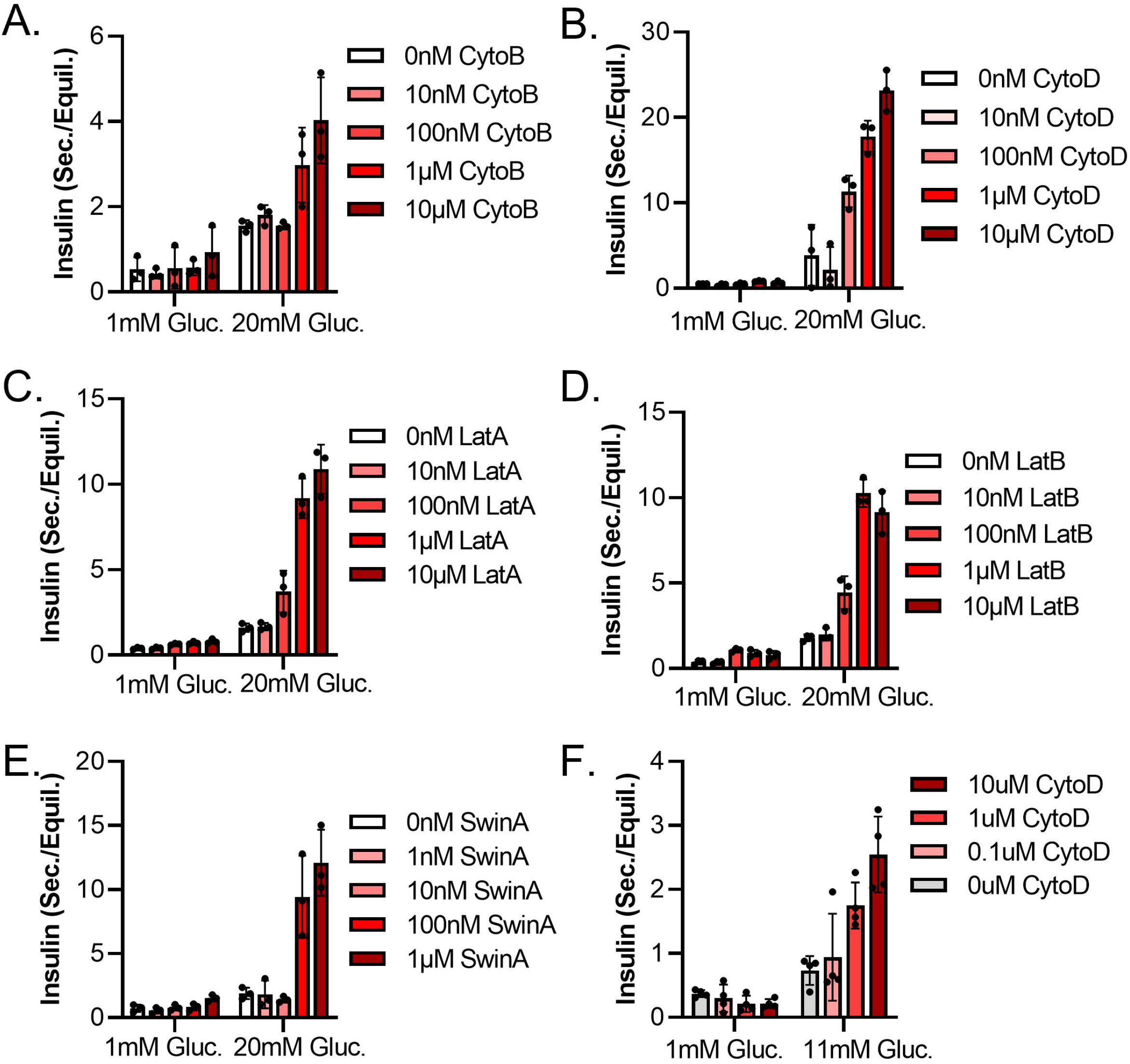
Actin filament disruptors enhance GSIS at all concentrations. (A-E) Insulin secretion assay from MIN6 cells. Cells were equilibrated for an hour in KRBH + 2.8 mM glucose (“Equil.”), pre-treated in KRBH + 2.8 mM glucose + the indicated inhibitor concentration, then treated with high (20 mM) or low (1 mM) glucose with the indicated amounts of inhibitor for one hour (“Sec.”). Points represent the amount of insulin in the “Sec.” sample of a given well divided by the amount of insulin in the “Equil.” sample from the same well. Three wells were tested per condition, with the insulin from each measured in technical duplicate. Each point represents the average of the technical duplicates from a single well. Bar height represents the mean of the three wells; error bars the standard deviation. Experiments were performed three times; a representative replicate of each is shown. (F) Insulin secretion assay from mouse islets. 4 islets per well; 4 wells per condition. Otherwise as above. Inhibitors are the actin filament disruptors cytochalasin B (CytoB), cytochalasin D (CytoD), latrunculin A (LatA), latrunculin B (LatB), and swinholide A (SwinA).

**Figure 4.**
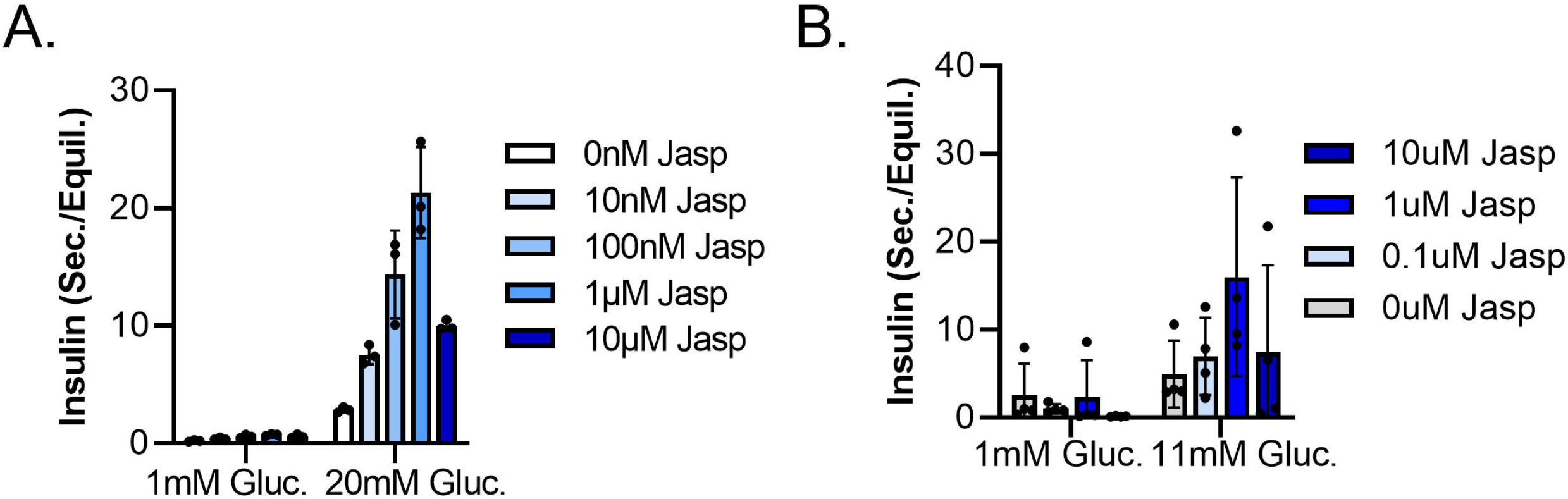
Actin filament-supporting drug also enhances GSIS at all concentrations. Insulin secretion assays from (A) MIN6 cells and (B) moues islet. Done exactly as in Figure 3. Jasplakinolide (Jasp).

### Actin cortex state does not explain insulin secretion phenotype

To determine if these drugs were acting by some shared effect on the cortical actin architecture, we again employed SEM on membrane-stripped mouse islets pre-treated with the actin depolymerizers cytochalasin D and latrunculin B, as well as the actin filament enhancer jasplakinolide (Fig. 5). Without drug, the actin cortex provides dense cover across the cell surface. Cytochalasin D or latrunculin B treatment dramatically strip away the topmost actin layer, revealing a network of thicker filaments and vesicles beneath. In contrast, after jasplakinolide treatment the cortical cytoskeleton is at least as dense as without drug. Thus, after drug treatment the cortical cytoskeleton state is discordant with the insulin secretion phenotype, suggesting these compounds enhance insulin secretion independent of the cortical cytoskeleton density.

**Figure 5.**
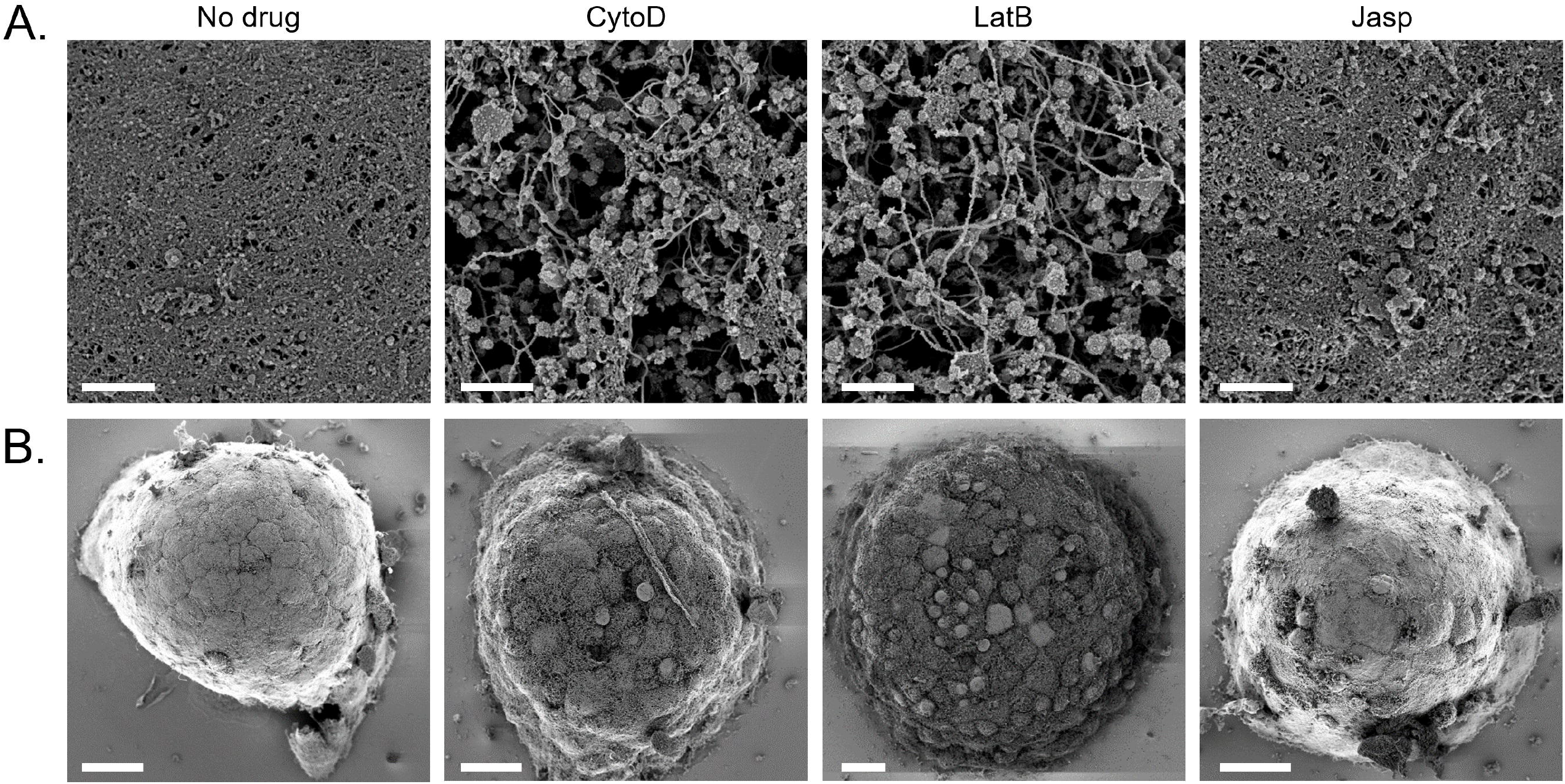
Cytoskeleton-modifying drugs that enhance GSIS have opposite effects on the cortical cytoskeleton. (A) Mouse islets were treated for 30 min. with KRBH plus 2.8 mM glucose (KRBH), or the same plus 1 μM cytochalasin D (CytoD), 1 μM latrunculin B (LatB), or 1 μM jasplakinolide (Jasp). Islets were then fixed, membranes extracted, and surfaces imaged by SEM as in Figure 2. Images are shown at 12,000x zoom. Scale bars represent 1 μm (B) Zoomed out images of islets from the same samples. Scale bars represent 20 μm. Experiment was performed twice with similar results; representative images are shown. More and uncropped images are shown in Fig. S3.

## Discussion

We set out to investigate the relationship between the cortical actin cytoskeleton and insulin secretion, testing a variety of compounds, concentrations, and treatment times in an attempt to unify apparently conflicting reports in the literature. We find that pro-secretory stimuli have little effect on the cortical actin network, both at a broad level across the cell, and at high-resolution. Further, we find drugs that either disrupt or enhance actin filament formation enhance GSIS, by a mechanism that is not directly tied to cortical cytoskeleton density. While these data suggest that the drugs are not affecting insulin secretion directly via the cortical cytoskeleton, the mechanism of action remains to be elucidated.

Our finding that stimulation causes little-to-no change in the actin cytoskeleton matches a recent study of dispersed mouse islets [9], but stands in contrast to several past studies [4–6,8]. We have attempted to recapitulate the conditions used in these studies, matching the treatment conditions and times to the extent possible. Perhaps some condition we were unable to account for – e.g. mouse/cell line genetics, cell density, or some unreported element of the media or substrate – explains the difference between our findings and others. However, in our hands, cells and islets secrete insulin in response to glucose or KCl without substantially degrading their actin cytoskeletons, suggesting the actin degradation that other groups have reported is not required for insulin secretion. We find pre-treatment of MIN6 cells or mouse islets with filament disruptors or enhancers potentiates GSIS as previously shown [11]. We extend this previous work by showing that this phenomenon is not unique to a particular actin-modifying compound or concentration, but is the case for several compounds in a concentration-dependent manner.

Why should these two classes of compounds with seemingly opposite effects on actin have the same effect on GSIS? We see two possibilities. (1) The two drugs act on different processes; perhaps actin disruptors act by perturbing cortical actin to allow granules greater access to secretory sites, while filament enhancers act by some unrelated (and yet unknown) mechanism. (2) The two drugs act on a single process, both influencing GSIS by freezing the dynamic turnover of actin filaments. In the second case, we might expect inhibition of actin-related processes (actomyosin motors, endocytosis, filament branching, et al.) to phenocopy filament disruption. However, inhibition of Arp2/3 (actin branching) [9], Cdc42 (involved in actin rearrangements) [12], FAK (signals through focal adhesions) [13], and myosin II (primary actomyosin motor) [14] have all been shown to decrease GSIS in *ex vivo* islets or primary β cells, consistent with these proteins being involved in the normal secretion process. In contrast, the formin inhibitor SMIFH2 (which also inhibits myosins) [9] and an inhibitor of Rac1 slightly enhance GSIS from moue islets, while the RhoA inhibitor rhosin dramatically enhances insulin secretion both at low and high glucose [12]. Of course, these references encompass a variety of β cell models, as well as drug concentrations and treatment timing, which limits our ability to draw a unified model of this pathway.

In summary, our data are not consistent with the longstanding model that cortical actin restricts secretion of insulin granules. Clearly, the actin cytoskeleton is substantially involved in the mechanism of insulin secretion, as even short-term perturbation of actin filaments or actin-related proteins can have a dramatic effect on insulin secretion. However, the nature of the cytoskeleton’s involvement remains unclear.

## Acknowledgements

We thank Silvia Jansen and David Kast for sharing their thoughts to shape this project, Sanja Sviben for technical support with the scanning electron microscopy, and all members of the Piston lab for supporting this project by sharing their thoughts and expertise.

## Methods

### Cell culture

MIN6 cells [15] were cultured in high glucose DMEM (Gibco, #11965092) supplemented with 10% fetal bovine serum (FBS), 50 μM β-mercaptoethanol, and 100 U/mL penicillin-streptomycin (Gibco, #15140122). Over the course of use, cells were monitored at least weekly to ensure they still retained glucose-stimulated insulin secretion.

### Rodent care and islet isolation

Rodent care was performed under the auspices of the Washington University Institutional Animal Care and Use Committee (AALAC accreditation # D16-00245). C57BL/6J mice were acquired from Jackson Laboratories (Strain #000664) or bred on-site and had access to water and chow *ad libitum*. Islets were isolated from both male and female mice less than 12 months old. Under anesthesia (100 mg/kg ketamine; 20 mg/kg xylazine) pancreata were surgically removed, then dissociated by shaking at 34 °C in HBSS (Gibco, #14025092) plus 0.75 mg/mL collagenase (Roche, #11213873001, dissolved at 12 mg/mL in HBSS without calcium – Gibco, #14175095) for 8 minutes, and washed three times in HBSS. Islets were picked manually under a stereomicroscope and allowed to recover overnight in RPMI 1640 (Gibco, #11879020) supplemented with 10% FBS, 11 mM glucose, and 100 U/mL penicillin-streptomycin.

### Immunofluorescence

For immunofluorescence MIN6 cells were seeded onto uncoated #1.5 coverslips (Electron Microscopy Sciences, #72230-01), while islets were seeded onto the same coverslips coated with recombinant human laminin-521 (Gibco, #A29248, 0.5 μg/cm ^2^). Cells/islets were allowed to attach overnight, rinsed three times in KRBH (128.8 mM NaCl, 4.8 mM KCl, 1.2 mM KH_2_PO_4_, 1.2 mM MgSO_4_, 2.5 mM CaCl_2_, 10 mM HEPES, 5 mM NaHCO_3_, 0.1% (w/v) bovine serum albumin (BSA), pH 7.4) plus 2.8 mM glucose, equilibrated in KRBH plus 2.8 mM glucose for one hour, treated in KRBH with the indicated amount of glucose (11 mM for islets, 20 mM for MIN6 cells) and KCl (0 mM or 30 mM) for the indicated amount of time (2-15 minutes). At the end of the treatment time, samples were fixed for 30 minutes in 2% paraformaldehyde (diluted from 16% solution, Thermo Fisher Scientific, #28906) in phosphate buffered saline (PBS, diluted from 10X, Corning, #46-013-CM). Fixed samples were rinsed three times in PBS, permeabilized in 0.1% Triton X-100 in PBS for 5 minutes, rinsed three times in BPS, and blocked for an hour with 2% BSA in PBS. After blocking, samples were stained with Phalloidin-AlexaFluor488 (Molecular Probes, #A12379) diluted in blocking buffer at 165 nM per the manufacturer’s recommendation. Samples were then washed three times with PBS, then coverslips inverted onto a drop of ProLong Glass Antifade Mountant (Invitrogen, #P36980), and allowed to cure for 48 hours before imaging.

All imaging was done on a Zeiss LSM880 laser-scanning confocal microscope using a Fluar 40x oil objective, numerical aperture 1.3, and a 32-PMT spectral detector array set to collect light from 495 nm to 630 nm. Images were taken at 1024 × 1024 pixels, a pixel dwell time of 1.03 μs, 2x line averaging, and a 2x zoom resulting in 0.1 μm per pixel. Images were processed in FIJI (ImageJ 1.53t), and graphed using GraphPad Prism (9.5).

### F-:G-actin assessment

To assess the ratio F-:G-actin, MIN6 cells were plated in a 24-well plate and allowed to adhere overnight. The next day, cells were equilibrated in KRBH plus 2.8 mM glucose for an hour, then treated for five minutes with KRBH + 1 mM glucose, KRBH + 20 mM glucose, or KRBH + 20 mM glucose + 30 mM KCl. Controls were treated for 30 minutes with KRBH + 2.8 mM glucose + 1 μM latrunculin B (Calbiochem, #428020) or 1 μM jasplakinolide (Millipore Sigma, #J4580). All samples were processed using the “G-actin / F-actin In Vivo Assay Kit” (Cytoskeleton Inc., #BK037) which involves lysing cells in a filament-supporting buffer, pelleting filaments by ultracentrifugation (100,000g; 1 hour), de-polymerizing filaments on ice, separating all by SDS-PAGE, and blotting for β-actin (buffers and antibody provided as part of kit). Secondary antibody was goat anti-mouse IRDye 800CW (LI-COR, #926-32210, used at 1:10,000). Blots were imaged on a LI-COR Odyssey M, and bands analyzed with Image Studio Lite (5.2.5).

### Scanning electron microscopy

Cells and islets were attached to coverslips and equilibrated in KRBH plus 2.8 mM glucose as in “Immunofluorescence” above. For images shown in Figure 2, following equilibration samples were treated with KRBH plus 2.8 mM glucose, KRBH plus high (20 mM for MIN6, 11 mM for mouse islets) glucose, or KRBH plus high glucose and 30 mM KCl for 5 minutes. For those shown in Figure 4, samples were treated for 30 minutes with KRBH plus 2.8 mM glucose plus 1 μM cytochalasin D, 1 μM latrunculin B, 1 μM jasplakinolide, or an equal volume of the solvent DMSO (1 μL in 1 mL).

In all cases, cells were then quickly rinsed in warm HBSS, fixed and extracted in PEM buffer (100 mM PIPES, pH 6.9; 1 mM EGTA; 1 mM MgCl_2_) plus 2% Triton X-100 and 0.25% glutaraldehyde (prepared fresh the day of experiment; a new ampule of glutaraldehyde was used each time) for 5 minutes with gentle shaking, followed by 10 minutes in PEM buffer + 2% Triton X-100 + 1% (w/v) CHAPS with gentle shaking. Samples were then rinsed three times in PEM buffer, and fixed in 0.15 M cacodylate buffer, pH 7.4 with 2% glutaraldehyde at 4 °C overnight.

Following fixation, coverslips were prepared for electron microscopy by treatment with 0.1% tannic acid for 20 minutes at room temperature, four rinses for five minutes each in water, 20 minutes in 0.2% aqueous uranyl acetate, three rinses for 10 minutes each in water, followed by a graded ethanol dehydration series with 5 minutes per step: 10%, 20%, 30%, 50%, 70%, 90%, 100%, 100%, 100%. Once dehydrated, samples were loaded into a critical point drier (Leica EM CPD 300) set to perform 12 CO_2_ exchanges at the slowest speed. Samples were mounted on aluminum stubs with carbon adhesive tabs and coated with 5 nm each of carbon and iridium (Leica ACE 600). Prepared samples were imaged on a Zeiss Merlin FE-SEM using an InLens detector at 1.2 – 1.5 kV and 400 – 500 pA depending on the sample. Images were scanned at 1024 × 768 pixels, resulting in 11.16 nm per pixel at the 10,000X zoom shown in Figure 2.

### Insulin secretion assay

For glucose-stimulated insulin secretion assays, MIN6 cells were plated in tissue-culture-treated 96-well plates and allowed to adhere overnight; mouse islets were plated on laminin-coated (0.5 μg/cm^2^, as above) half-area 96-well plates, 4 islets per well and allowed to adhere overnight. On the day of experiment, samples were washed three times in KRBH plus 2.8 mM glucose, then equilibrated in KRBH plus 2.8 mM glucose for one hour. The equilibration sample was saved, and replaced with a pre-treatment buffer containing KRBH plus 2.8 mM glucose plus inhibitors at the indicated concentrations (see below, “Inhibitors”) for 30 minutes. Cells were then moved to a “secretion buffer” of KRBH plus high glucose (20 mM for MIN6; 11 mM for mouse islets) plus inhibitors at the indicated concentrations for one hour. Secretion buffer was saved. Secretion and equilibration samples were spun at 1000 g for 3 min to remove any cellular material, and diluted as-needed in KRBH to be within the range of detection of our insulin assay. Insulin in each sample was measured in technical duplicate using the Lumit Insulin Immunoassay (Promega, #CS3037A01, provided through Promega’s early access program), with luminescence measured on a multimodal plate reader (Cytation5, BioTek). For MIN6 assays, three wells were assayed per condition; for islets (which have more variable size, composition, and signal) four wells were assayed per condition. Data were graphed using Graphpad Prism (9.5) and are shown throughout with a dot representing the insulin content in a given well’s secretion buffer over the same well’s equilibration buffer. Bar height represents the mean of the 3-4 wells measured in that condition. Error bars represent standard deviation.

### Inhibitors

All inhibitors described here were dissolved in DMSO and stored as aliquots at -70 °C until time of use. Latrunculin A (Millipore-Sigma, #L5163), latrunculin B (Calbiochem, #428020), cytochalasin B (Millipore-Sigma, #C6762), cytochalasin D (Millipore-Sigma, #C8273), swinholide A (Millipore-Sigma, #574776), jasplakinolide (Millipore-Sigma, #J4580). For glucose-stimulated insulin secretion assays, inhibitors were diluted into KRBH to the highest experimental concentration, then diluted with 1:10 serial dilutions in KRBH to make the remaining concentrations.

## References

1. Chugh P, Paluch EK. The actin cortex at a glance. J Cell Sci. 2018;131: jcs186254. doi:10.1242/jcs.186254

2. Li P, Bademosi AT, Luo J, Meunier FA. Actin Remodeling in Regulated Exocytosis: Toward a Mesoscopic View. Trends in Cell Biology. 2018;28: 685–697. doi:10.1016/j.tcb.2018.04.004

3. Nightingale TD, Cutler DF, Cramer LP. Actin coats and rings promote regulated exocytosis. Trends in Cell Biology. 2012;22: 329–337. doi:10.1016/j.tcb.2012.03.003

4. Asahara S, Shibutani Y, Teruyama K, Inoue HY, Kawada Y, Etoh H, et al. Ras-related C3 botulinum toxin substrate 1 (RAC1) regulates glucose-stimulated insulin secretion via modulation of F-actin. Diabetologia. 2013;56: 1088–1097. doi:10.1007/s00125-013-2849-5

5. Kalwat MA, Yoder SM, Wang Z, Thurmond DC. A p21-activated kinase (PAK1) signaling cascade coordinately regulates F-actin remodeling and insulin granule exocytosis in pancreatic β cells. Biochem Pharmacol. 2013;85: 808–816. doi:10.1016/j.bcp.2012.12.003

6. Bowe JE, Chander A, Liu B, Persaud SJ, Jones PM. The permissive effects of glucose on receptor-operated potentiation of insulin secretion from mouse islets: a role for ERK1/2 activation and cytoskeletal remodelling. Diabetologia. 2013;56: 783–791. doi:10.1007/s00125-012-2828-2

7. Naumann H, Rathjen T, Poy MN, Spagnoli FM. The RhoGAP Stard13 controls insulin secretion through F-actin remodeling. Mol Metab. 2018;8: 96–105. doi:10.1016/j.molmet.2017.12.013

8. Wang B, Lin H, Li X, Lu W, Kim JB, Xu A, et al. The adaptor protein APPL2 controls glucose-stimulated insulin secretion via F-actin remodeling in pancreatic β-cells. Proc Natl Acad Sci U S A. 2020;117: 28307–28315. doi:10.1073/pnas.2016997117

9. Ma W, Chang J, Tong J, Ho U, Yau B, Kebede MA, et al. Arp2/3 nucleates F-actin coating of fusing insulin granules in pancreatic β cells to control insulin secretion. J Cell Sci. 2020;133: jcs236794. doi:10.1242/jcs.236794

10. Orci L, Gabbay KH, Malaisse WJ. Pancreatic beta-cell web: its possible role in insulin secretion. Science. 1972;175: 1128–1130. doi:10.1126/science.175.4026.1128

11. Mourad NI, Nenquin M, Henquin J-C. Amplification of insulin secretion by acetylcholine or phorbol ester is independent of β-cell microfilaments and distinct from metabolic amplification. Mol Cell Endocrinol. 2013;367: 11–20. doi:10.1016/j.mce.2012.12.002

12. Ng XW, Chung YH, Asadi F, Kong C, Ustione A, Piston DW. RhoA as a signaling hub controlling glucagon secretion from pancreatic alpha-cells. 2021.

13. Jevon D, Deng K, Hallahan N, Kumar K, Tong J, Gan WJ, et al. Local activation of focal adhesion kinase orchestrates the positioning of presynaptic scaffold proteins and Ca2+ signalling to control glucose dependent insulin secretion. Jahn R, editor. eLife. 2022;11: e76262. doi:10.7554/eLife.76262

14. Arous C, Rondas D, Halban PA. Non-muscle myosin IIA is involved in focal adhesion and actin remodelling controlling glucose-stimulated insulin secretion. Diabetologia. 2013;56: 792–802. doi:10.1007/s00125-012-2800-1

15. Miyazaki J, Araki K, Yamato E, Ikegami H, Asano T, Shibasaki Y, et al. Establishment of a pancreatic beta cell line that retains glucose-inducible insulin secretion: special reference to expression of glucose transporter isoforms. Endocrinology. 1990;127: 126–132. doi:10.1210/endo-127-1-126

