## Supplemental Figures for "Disrupting actin filaments enhances glucose-stimulated insulin secretion independent of the cortical actin cytoskeleton"

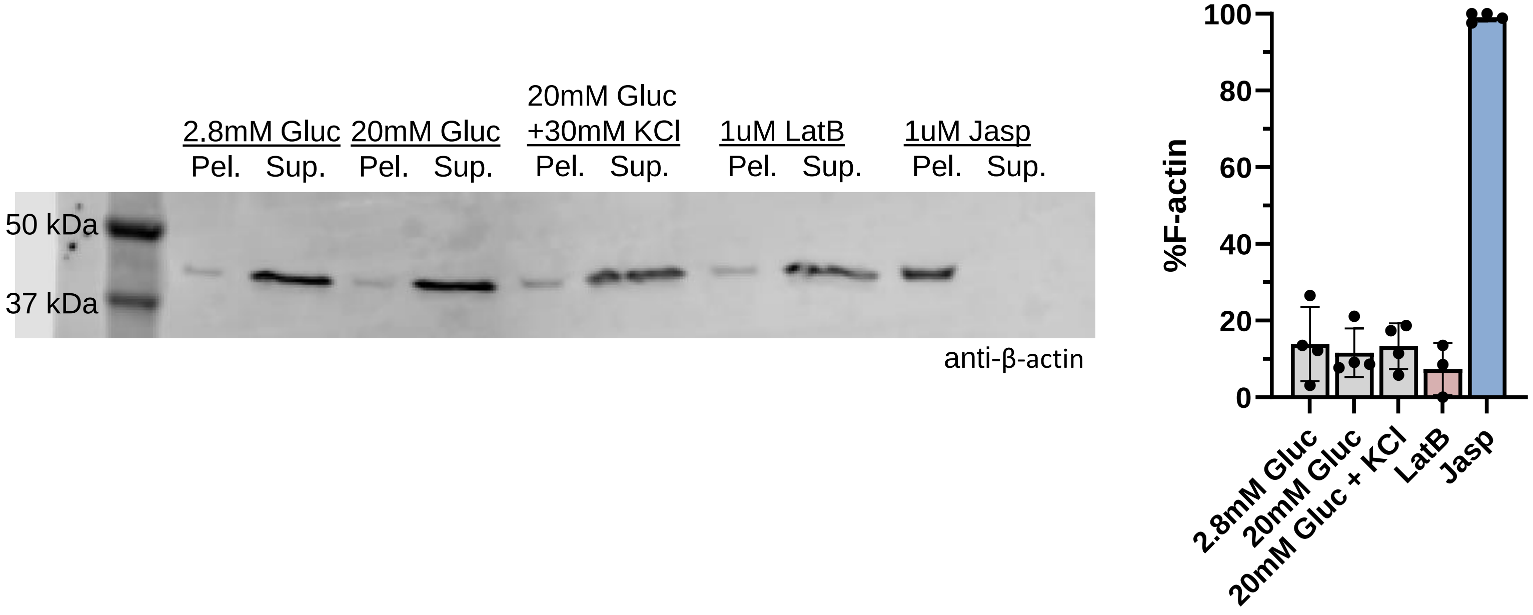


**Figure S1. Stimulating MIN6 cells does not substantially change ratio F-:G-actin** – MIN6 cells were stimulated with low glucose (1 mM), high glucose (20 mM), or high glucose plus 30 mM KCl for 5 minutes, then actin filaments separated by ultracentrifugation – filaments fall to the pellet (“Pel.”) while unfilamented actin remains in the supernatant (“Sup.”) – and assessed by western blot. 30 minutes treatments with latrunculin B (depolymerizes filaments) and jasplakinolide (polymerizes filaments) serve as controls. Experiment was performed four times. A representative experiment is shown (left), while all experiments are quantified at right. Bars represent standard deviation of the measurements.

Figure S2: Zoomed out photos from Fig. 2

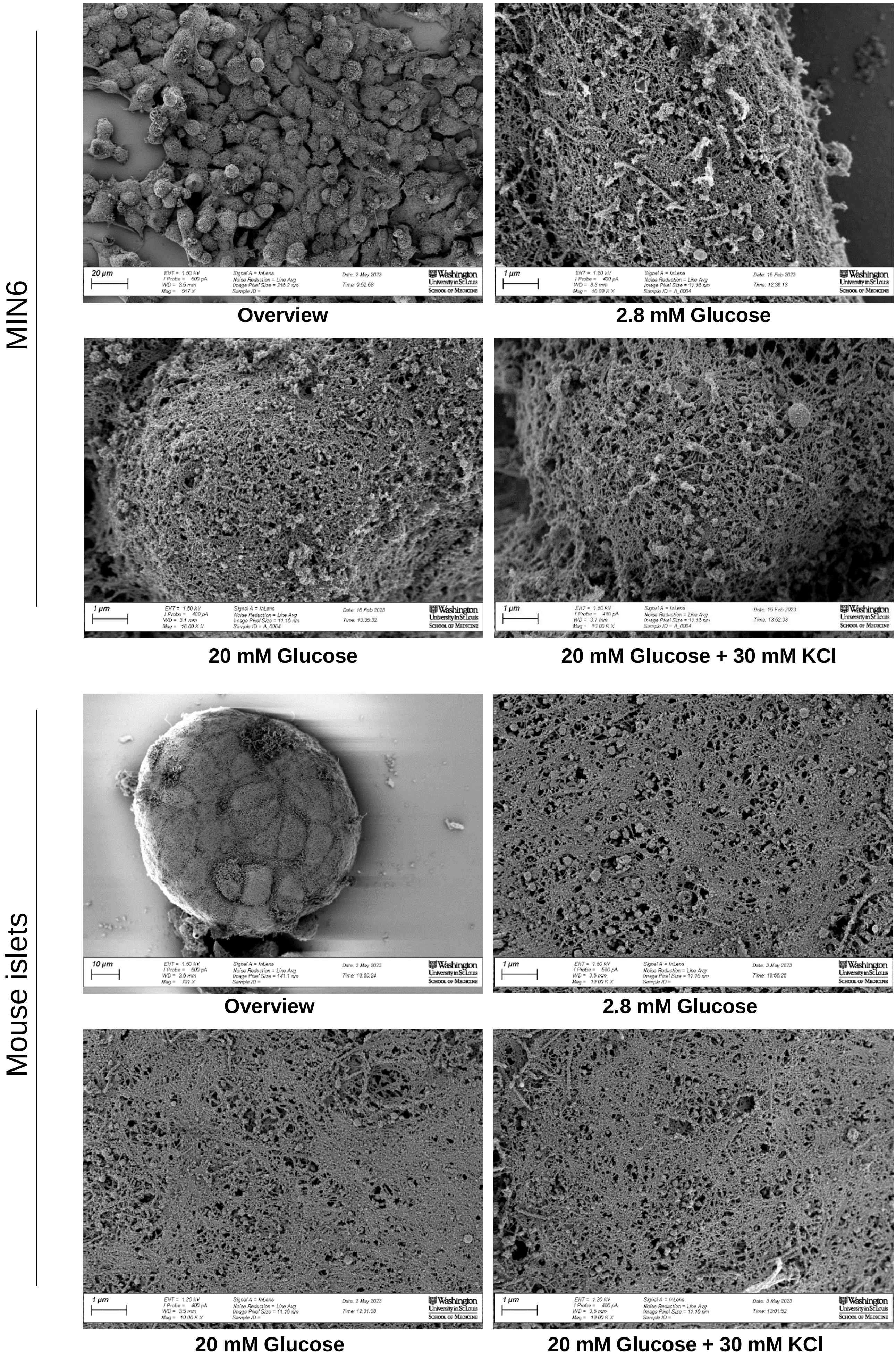

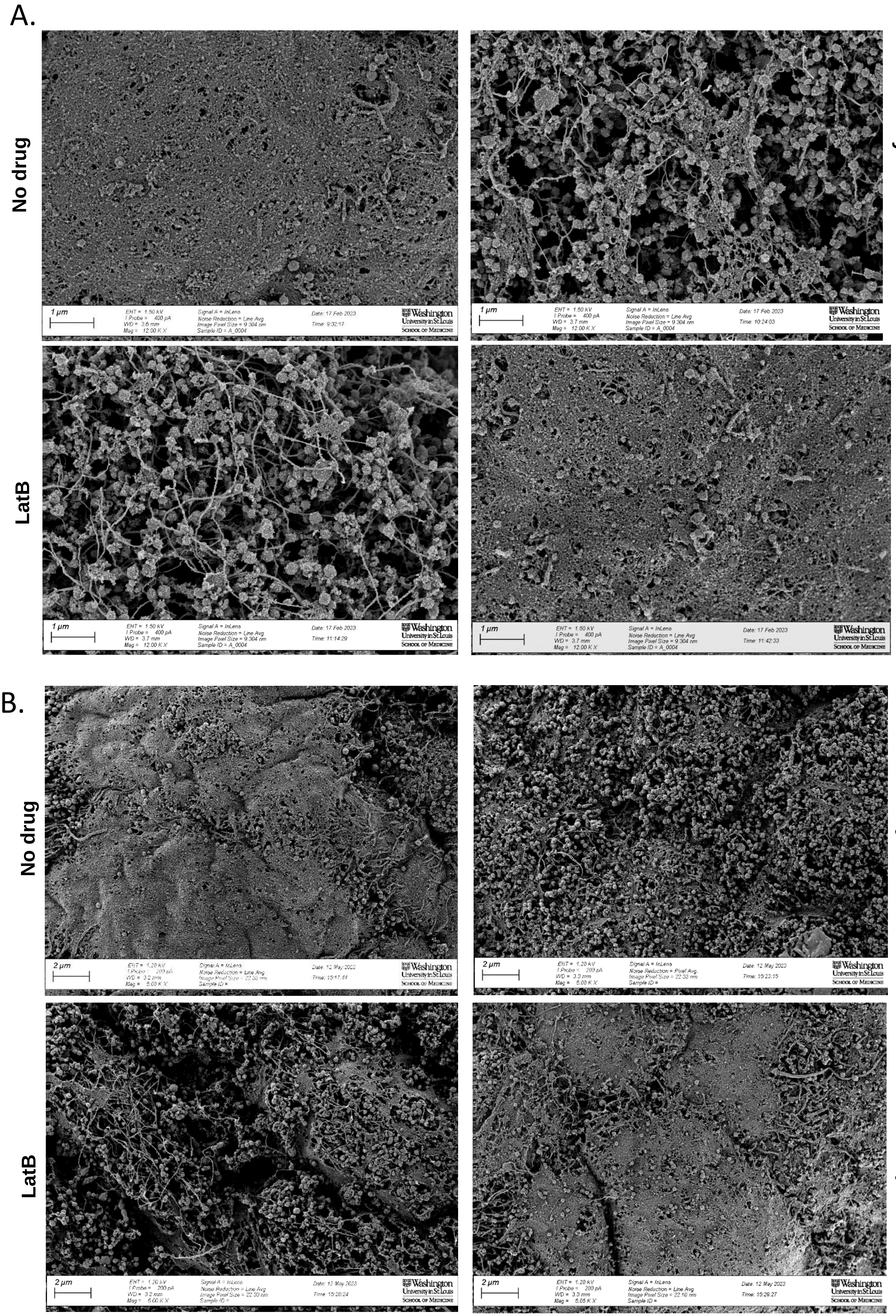


**Figure S3. More and uncropped photos from Fig. 5 –** (A) Uncropped photos from Fig. 5A, (B) An additional view from each samples at 5,000x
